# Frequency-dependent Distribution of Different Subtypes of Spiral Ganglion Neurons in the Cochlea and the Changes During Aging

**DOI:** 10.1101/2023.05.19.541526

**Authors:** Meijian Wang, Shengyin Lin, Ruili Xie

## Abstract

Sound information is transmitted from the cochlea to the brain mainly by type I spiral ganglion neurons (SGNs), which consist of different subtypes with distinct physiological properties and selective expression of molecular markers. It remains unclear how these SGN subtypes distribute along the tonotopic axis, and how the distribution pattern changes during aging that might underlie age-related hearing loss (ARHL). We investigated these questions using immunohistochemistry in three age groups of CBA/CaJ mice of either sex, including 2-5 months (young), 17-19 months (middle-age), and 28-32 months (old). Mouse cochleae were cryo-sectioned and triple-stained using antibodies against Tuj1, calretinin (CR) and calbindin (CB), which are reportedly expressed in all type I, subtype I_a_, and subtype I_b_ SGNs, respectively. Labeled SGNs were classified into four groups based on the expression pattern of stained markers, including CR^+^ (subtype I_a_), CB^+^ (subtype I_b_), CR^+^CB^+^ (dual-labeled I_a_/I_b_), and CR^-^CB^-^ (subtype I_c_) neurons. The distribution of these SGN groups was analyzed in the apex, middle, and base regions of the cochleae. It showed that the prevalence of subtype I_a_, I_b_ and dual-labeled I_a_/I_b_ SGNs are high in the apex and low in the base. In contrast, the distribution pattern is reversed in I_c_ SGNs. Such frequency-dependent distribution is largely maintained during aging except for a preferential reduction of I_c_ SGNs, especially in the base. These findings suggest that sound processing of different frequencies involves distinct combinations of SGN subtypes, and the age-dependent loss of I_c_ SGNs in the base may especially impact high-frequency hearing during ARHL.

## Introduction

Type I spiral ganglion neurons (SGN) make up 95% of all SGNs and are the main cells to transmit sound information from the sensory hair cells to the brain^1, 2^. These neurons show diverse physiological properties and were traditionally classified into different subtypes based on their spontaneous firing rate and threshold to sound^3-6^. Specifically, type I SGNs are composed of high spontaneous rate (HSR) neurons with low sound threshold, medium spontaneous rate (MSR) neurons with medium sound threshold, and low spontaneous rate (LSR) neurons with high sound threshold^3^. LSR SGNs show a much broader dynamic range in encoding sound intensity than HSRs^4, 7^, and are believed to be important in detecting transient^8^ and loud signals in noise^9, 10^. Morphologically, these SGNs differ in dendritic fiber caliber^2, 11^, cochlear synaptic location and structure on inner hair cells (IHC)^2, 12^, and their central projections and synaptic structure in the cochlear nucleus^13-15^. Particularly, HSR neurons have thick fiber diameter and synapse onto the modiolar side of IHCs, while LSR neurons have thin fiber diameter and synapse onto the pillar side of IHCs^2^. Studies also showed that different subtypes of SGNs are not evenly distributed along the frequency axis^5, 16-18^, and are differentially altered under various pathological conditions including aging^18-20^. However, the comprehensive distribution pattern of these SGNs across tonotopy and the age-related changes have not been well-characterized in detail.

Recent studies using single-cell RNA sequencing technique identified various molecular markers that define three subtypes of type I SGNs^18, 21, 22^. Of particular interest, calretinin (CR) is expressed in subtype I_a_ SGNs, calbindin (CB) is expressed in subtype I_b_ SGNs, whereas Pou4f1 is expressed in subtype I_c_ SGNs. Additional studies based on synaptic location^18, 21, 23, 24^ and physiological property^25^ showed that the molecularly identified subtype I_a_, I_b_ and I_c_ SGNs roughly correspond to traditionally classified HSR, MSR and LSR SGNs. These findings make studies possible to conveniently identify SGN cell types and investigate the comprehensive distribution pattern of different SGN subtypes across tonotopic axis.

In this study, we used immunohistochemistry to identify various SGN subtypes in cryo-sectioned cochlea by simultaneously staining three molecular markers of Tuj1, CR, and CB^18, 21, 26^. Four groups of type I SGNs were classified based on the expression pattern of these markers, including CR^+^ (subtype I_a_), CB^+^ (subtype I_b_), CR^+^CB^+^ (dual-labeled I_a_/I_b_), and CR^-^CB^-^ (subtype I_c_) neurons. We found that all SGN subtypes showed gradient distribution across frequencies, with the proportions of subtype I_a_, I_b_, and dual-labeled I_a_/I_b_ SGNs decreased from the apex to the base regions of the cochlea, and the proportion of subtype I_c_ SGNs reciprocally increased. The overall SGN density decreased with age, reflecting age-dependent SGN loss. Furthermore, the cell loss preferentially impacted subtype I_c_ SGNs in the base region. The frequency-dependent distribution of SGN subtypes indicates that subtype I_a_ and I_b_ SGNs contribute more to the processing of low frequency sound, and subtype I_c_ SGNs play more roles in the processing of high frequency sound. During aging, preferential loss of I_c_ SGNs in the base region may uniquely contribute to high frequency hearing deficit during ARHL.

## Methods

Animal experiments were conducted under the guidelines of the protocol #2018A00000055, approved by the Institutional Animal Care and Use Committee of The Ohio State University.

CBA/CaJ mice were acquired from the Jackson Laboratory and maintained at the animal facility at The Ohio State University. Mice of either sex were used at three age groups, including 2-5 months (young), 17-19 months (middle age), and 28-32 months (old), which represent normal hearing, hidden hearing loss, and severe ARHL, respectively^27^. As previously reported^27, 28^, hearing status of these mice was verified by recording auditory brainstem response under anesthesia after an I.P. injection of ketamine (100 mg/kg) and xylazine (10 mg/kg). Mice were decapitated to retrieve the cochleae, which were fixed in PBS with 4% paraformaldehyde for overnight at 4 °C and de-calcified in 0.12 M EDTA in 0.1 M PBS for 2-3 days. Cochleae were then cryo-protected in 30% sucrose in PBS, embedded in Cryo-Gel (Cat. #:475237; Instrumedics Inc.) and sectioned along the modiolus axis using a cryostat slicer (Leica CM3050 S, Leica Biosystems), at the thickness of 20 μm.

Cochlea sections were triple-stained using primary antibodies against Tuj1 (rabbit anti-Tuj1, Cat# ab52623; Abcam), CR (guinea pig anti-CR, Cat# 214104; Synaptic Systems), and CB (chicken anti-CB, Cat# 214006; Synaptic Systems). Corresponding secondary antibodies were used including goat anti-guinea pig - Alexa Fluor 488 (Cat# A-11073; Invitrogen), goat anti-rabbit - Alexa Fluor 594 (Cat# A-11037; Invitrogen), goat anti-chicken - Alexa Fluor 647 (Cat# A-32933; Invitrogen), and goat anti-rabbit - Alexa Fluor 750 (Cat# A-21039; Invitrogen). Slices were mounted using DAPI Fluoromount-G mounting medium (Cat# 0100-20; Southern Biotech) to label cell nucleus. Stained cochleae were imaged using an Olympus FV3000 confocal microscope (Olympus). Z-stack images were acquired across 9 μm in depth at the step of 1.5 μm.

Image analysis used Imaris version 9.9 (Oxford Instruments). SGN cell bodies were reconstructed using the “Spot” tool based on the Tuj1 staining. SGN cell number was quantified. Subtype-specificity of each SGN was determined by the staining pattern of the above three molecular markers. Specifically, four subgroups of SGNs were quantified among all Tuj1-labelled type I SGNs in the apex, middle and basal portions of the spiral ganglia, including those that express CR (CR^+^), CB (CB^+^), both CR and CB (CR^+^CB^+^), and none of the two (CR^-^CB^-^). SGN density was calculated as the SGN number divided by the measured area of the ganglion region.

Statistical analysis was performed using Prism, version 6.0h (GraphPad Software Inc.). Both one-way and two-way ANOVA test were used to evaluate the effects of frequency region and age on the distribution of SGN subtypes. Data are presented as mean ± SD. Only one cochlea per mouse was used, and each reported data point represents an individual animal.

## Results

### Identification of SGN subtypes and their frequency-dependent distribution in young mice

We first investigated the distribution of SGN subtypes in young CBA/CaJ mice. As shown in Fig. 1A, cell bodies of type I SGNs were clearly labeled by Tuj1-staining^26^ in the apex, middle, and base regions of the spiral ganglia in a single cochlea section. Additional CR- and CB-staining labeled different subpopulations of SGNs, as visualized by different colors in the merged panel. To identify and quantify different SGNs, we reconstructed all three regions of the spiral ganglia using balls of different colors to represent the cell bodies of labeled SGNs. As shown in Fig. 1B, CR^+^ SGNs are represented as red balls among all Tuj1-labeled neurons in the left two columns (CR+Tuj1 overlap), CB^+^ SGNs are shown as green balls in the middle two columns (CB+Tuj1 overlap). There are relatively more red and green balls in the apex than the middle and base regions, revealing a frequency-dependent distribution of CR^+^ and CB^+^ SGNs. Overlap of all three stained markers (right two columns) showed that many SGNs were double-labeled by CR and CB, which we classified as CR^+^CB^+^ neurons (brown balls). Neurons without CR or CB labeling were classified as CR^-^CB^-^ SGNs, represented as blue balls in the right column, which showed clear increase in prevalence toward the base region.

**Figure 1.**
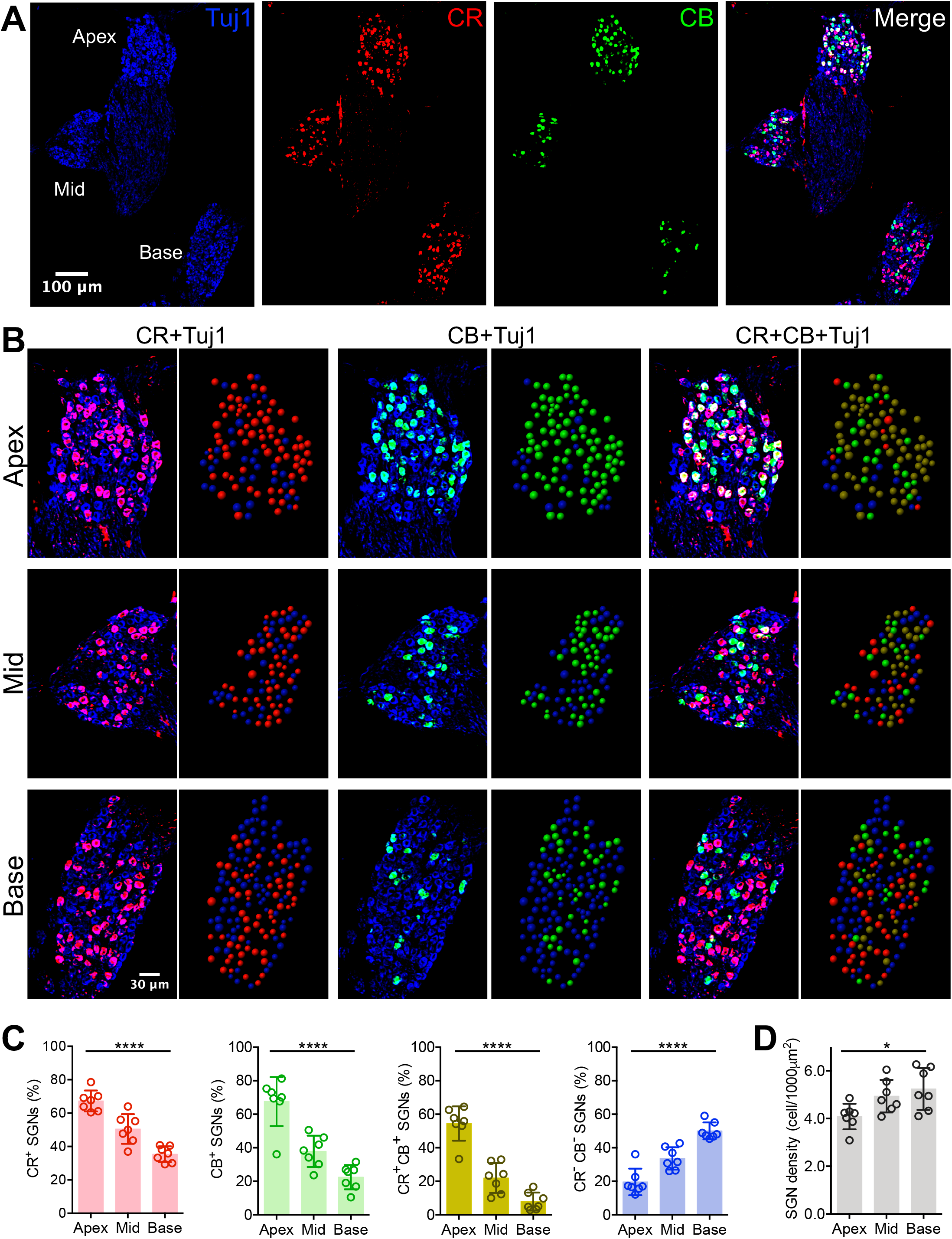
Frequency-dependent distribution of different subtypes of type I SGNs in young mice. (**A**) Cryo-section of an example cochlea stained with Tuj1, calretinin (CR) and calbindin (CB). Notice that CR and CB label different sub-populations of Tuj1^+^ SGNs and their distributions vary across different frequency regions. (**B**) Magnified view of (**A**) to show staining of SGNs in apex, middle and base regions. Color-coded balls on right panels represent identified SGNs. Red: CR^+^ SGNs; green: CB^+^ SGNs. Brown and blue in the rightmost panels represent CR^+^CB^+^ and CR^-^CB^-^ SGNs, respectively. (**C**) Percentage distribution of different subtypes of SGNs in three regions. (**D**) Average SGN cell density in three regions. One-way ANOVA: * P < 0.05; **** P < 0.0001.

We summarized the cellular prevalence of different SGNs in 7 young mice, and found that CR^+^, CB^+^ and CR^+^CB^+^ neurons were more abundant in the apex and fewer in the base; while the CR^-^CB^-^ neurons showed an opposite distribution pattern (Fig. 1C). On average, the prevalence of CR^+^ neurons was 67 ± 6% in the apex, 51 ± 9% in the middle, and 35 ± 5% in the base region of the spiral ganglia (one-way ANOVA: F_(2, 18)_ = 38.2; P < 0.0001). The prevalence of CB^+^ neurons was 68 ± 15% in the apex, 38 ± 9% in the middle, and 22 ± 7% in the base region of the spiral ganglia (one-way ANOVA: F_(2, 18)_ = 31.3; P < 0.0001). The prevalence of CR^+^CB^+^ neurons was 54 ± 10% in the apex, 22 ± 9% in the middle, and 8 ± 5% in the base region of the spiral ganglia (one-way ANOVA: F_(2, 18)_ = 56.1; P < 0.0001). In contrast, the prevalence of CR^-^CB^-^ neurons was 20 ± 8% in the apex, 34 ± 7% in the middle, and 50 ± 5% in the base regions of the spiral ganglia (one-way ANOVA: F_(2, 18)_ = 37.0; P < 0.0001). As a reference, the overall cell density of SGNs was 4.1 ± 0.5 cell/1000 μm^2^ in the apex, 4.9 ± 0.7 cell/1000 μm^2^ in the middle, and 5.2 ± 0.9 cell/1000 μm^2^ in the base regions of the spiral ganglia (Fig. 1D; one-way ANOVA: F_(2, 18)_ = 5.0; P = 0.019). The results suggest that sound encoding at different frequencies are achieved by different populations of SGN subtypes, in which CR^+^ and CB^+^ SGNs play a larger role in encoding low frequency sound and CR^-^CB^-^ SGNs play a larger role in encoding high frequency sound.

### Distribution of SGN subtypes changes with age

To investigate the age-dependent changes of SGN subtypes, we extended our study to include 5 middle-age and 7 old mice. Representative examples of cochlea staining are shown in Fig. 2, including the apex (Fig. 2A), middle (Fig. 2B) and base (Fig. 2C) regions of the spiral ganglia from all three age groups. While the prevalence of CR^+^ and CB^+^ neurons remained relatively steady during aging in the apex region (Fig. 2A), both were clearly increased with age in the middle (Fig. 2B) and base (Fig. 2C) regions. Consistently, the prevalence of double labeled CR^+^CB^+^ neurons (brown balls in merged panels) were also increased. In contrast, the prevalence of CR^-^CB^-^ neurons (blue balls in merged panels) were decreased with age in the middle and base regions.

**Figure 2.**
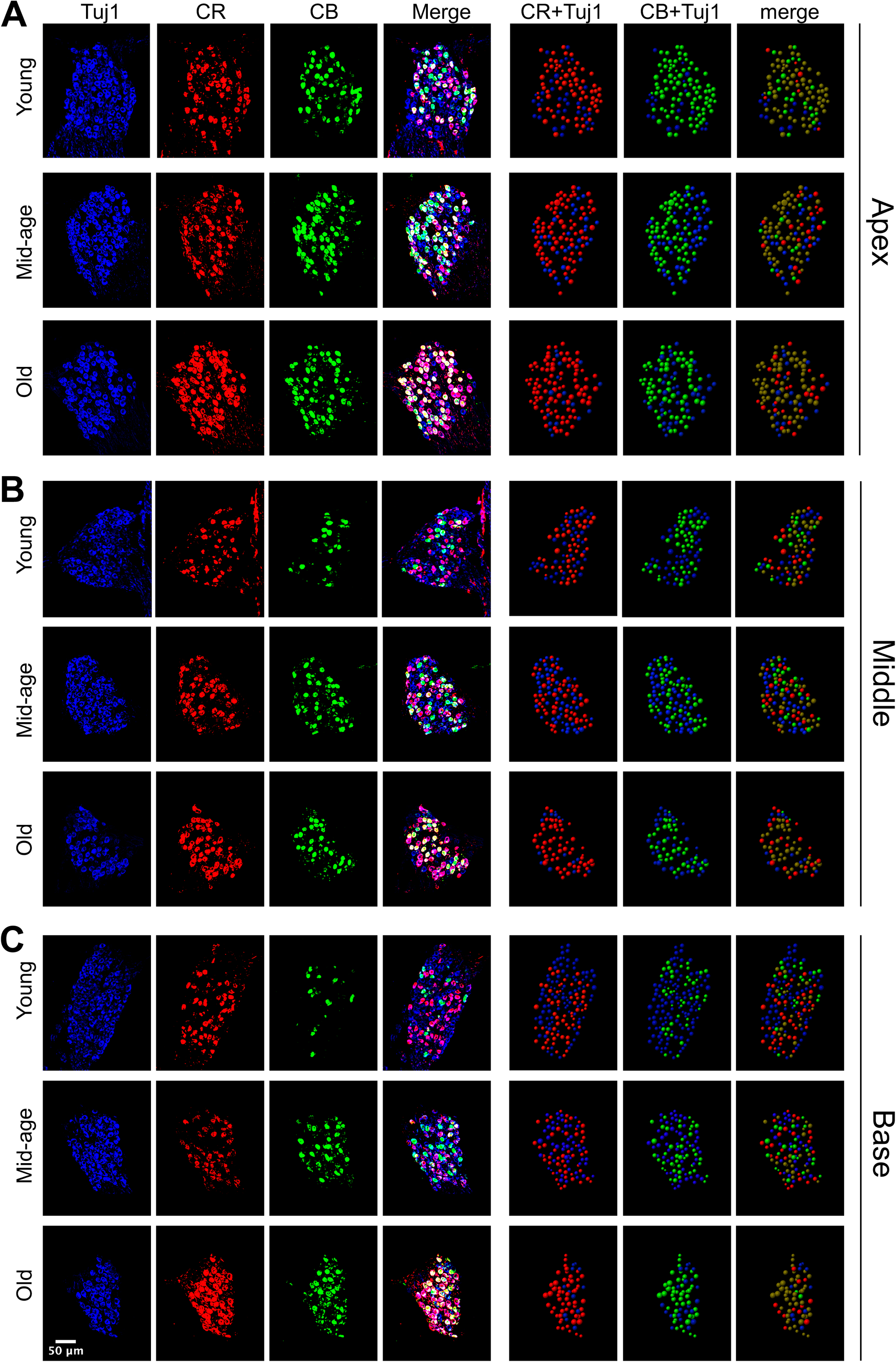
Age-dependent changes in the distribution of different subtypes of type I SGNs. (**A-C**) Staining of example cochleae from young, middle-age, and old mice across three frequency regions in the apex (**A**), middle (**B**), and base (**C**). Color-coded balls on right panels represent identified SGNs. Brown and blue balls in the rightmost column represent CR^+^CB^+^ and CR^-^CB^-^ SGNs, respectively.

Specifically, as shown in Fig. 3A, the prevalence of CR^+^ neurons in the apex region was 67 ± 6% in young, 74 ± 9% in middle-age, and 78 ± 4% in old mice (one-way ANOVA: F_(2, 16)_ = 4.7; P = 0.025); In the middle region, it was 51 ± 9% in young, 56 ± 3% in middle-age, and 72 ± 5% in old mice (one-way ANOVA: F_(2, 16)_ = 20.0; P < 0.0001); In the base region, it was 35 ± 5% in young, 49 ± 9% in middle-age, and 70 ± 9% in old mice (one-way ANOVA: F_(2, 16)_ = 36.8; P < 0.0001). Two-way ANOVA showed significant main effects of age and frequency region (age effect: F_(2, 48)_ = 55.4, P < 0.0001; frequency region effect: F_(2, 48)_ = 46.7, P < 0.0001; interaction: F_(4, 48)_ = 5.9, P = 0.0006). Similarly, the prevalence of CB^+^ neurons in the apex region was 68 ± 15% in young, 67 ± 18% in middle-age, and 71 ± 9% in old mice (one-way ANOVA: F_(2, 16)_ = 0.17; P = 0.847); In the middle region, it was 38 ± 9% in young, 52 ± 9% in middle-age, and 59 ± 7% in old mice (one-way ANOVA: F_(2, 16)_ = 10.8; P = 0.0011); In the base region, it was 22 ± 7% in young, 37 ± 15% in middle-age, and 55 ± 14% in old mice (one-way ANOVA: F_(2, 16)_ = 12.5; P = 0.0005). Two-way ANOVA showed significant main effects of age and frequency region (age effect: F_(2, 48)_ = 13.7, P < 0.0001; frequency region effect: F_(2, 48)_ = 32.5, P < 0.0001; interaction: F_(4, 48)_ = 2.8, P = 0.036). The prevalence of CR^+^CB^+^ neurons in the apex region was 54 ± 10% in young, 57 ± 18% in middle-age, and 63 ± 7% in old mice (one-way ANOVA: F_(2, 16)_ = 1.00; P = 0.390); In the middle region, it was 22 ± 9% in young, 38 ± 7% in middle-age, and 54 ± 6% in old mice (one-way ANOVA: F_(2, 16)_ = 31.2; P < 0.0001); In the base region, it was 8 ± 5% in young, 20 ± 8% in middle-age, and 45 ± 16% in old mice (one-way ANOVA: F_(2, 16)_ = 19.8; P < 0.0001). Two-way ANOVA showed significant main effects of age and frequency region (age effect: F_(2, 48)_ = 33.2, P < 0.0001; frequency region effect: F_(2, 48)_ = 50.4, P < 0.0001; interaction: F_(4, 48)_ = 3.9, P = 0.0085). Finally, the prevalence of CR^-^CB^-^ neurons in the apex region was 20 ± 8% in young, 16 ± 8% in middle-age, and 14 ± 5% in old mice (one-way ANOVA: F_(2, 16)_ = 1.04; P = 0.378); In the middle region, it was 34 ± 7% in young, 30 ± 4% in middle-age, and 23 ± 6% in old mice (one-way ANOVA: F_(2, 16)_ = 6.43; P = 0.0089); In the base region, it was 50 ± 5% in young, 35 ± 15% in middle-age, and 20 ± 6% in old mice (one-way ANOVA: F_(2, 16)_ = 18.6; P < 0.0001). Two-way ANOVA showed significant main effects of age and frequency region (age effect: F_(2, 48)_ = 22.4, P < 0.0001; frequency region effect: F_(2, 48)_ = 29.9, P < 0.0001; interaction: F_(4, 48)_ = 5.4, P = 0.0011).

**Figure 3.**
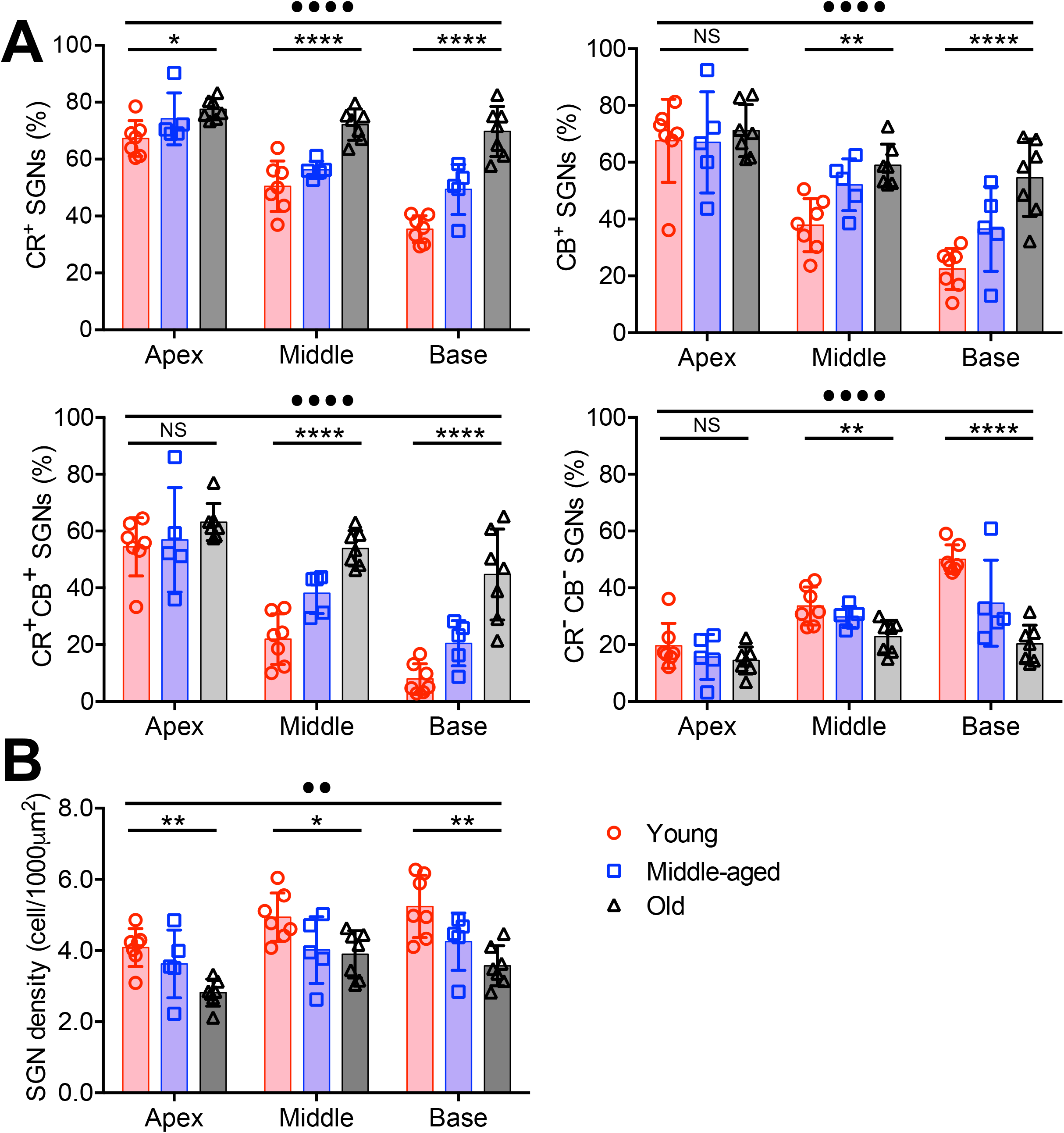
Summary distribution of different subtypes of type I SGNs during aging. (**A**) Summary distributions of CR^+^, CB^+^, CR^+^CB^+^, and CR^-^CB^-^ SGNs during aging in the apex, middle and base regions of the cochleae. (**B**) Changes in the average SGN cell density during aging across three frequency regions. One-way ANOVA: * P < 0.05; ** P < 0.01; **** P < 0.0001; NS: not significant. Two-way ANOVA: •• P < 0.01; •••• P < 0.0001.

Overall, SGN density was significantly decreased with age, reflecting the loss of SGNs during aging (Fig. 3B). In the apex region, the average SGN density was 4.1 ± 0.5 in young, 3.6 ± 1.0 in middle-age, and 2.8 ± 0.4 cell/1000 μm^2^ in old mice (one-way ANOVA: F_(2, 16)_ = 7.4; P = 0.0054); In the middle region, the average SGN density was 4.9 ± 0.7 in young, 4.0 ± 0.9 in middle-age, and 3.9 ± 0.7 cell/1000 μm^2^ in old mice (one-way ANOVA: F_(2, 16)_ = 3.9; P = 0.040); In the base region, the average SGN density was 5.2 ± 0.9 in young, 4.2 ± 0.8 in middle-age, and 3.6 ± 0.6 cell/1000 μm^2^ in old mice (one-way ANOVA: F_(2, 16)_ = 8.6; P = 0.0029). Two-way ANOVA showed significant main effects of age and frequency region (age effect: F_(2, 48)_ = 18.4, P < 0.0001; frequency region effect: F_(2, 48)_ = 8.1, P = 0.0010; interaction: F_(4, 48)_ = 0.65, P = 0.629).

The results showed that during aging, the overall number of SGNs gradually decreased in all three frequency regions due to cell loss (Fig. 3B). Furthermore, the age-dependent SGN loss is more prominent in CR^-^CB^-^ neurons, especially in the middle and base regions of the spiral ganglia (Fig. 3A). Accordingly, the prevalence of CR^+^ and CB^+^ are relatively increased, leading to altered composition of different SGN subtypes across different frequency regions of the cochlea. Given that different SGN subtypes have distinct physiological function in processing sound, such altered distribution during aging is expected to differentially impact hearing across different frequencies, especially contributing to the typical high frequency hearing loss in ARHL.

## Discussion

Using immunohistochemistry with three molecular markers, this study systematically characterized the distribution patterns of different subtypes of type I SGNs and their age-related changes in the cochlea across three frequency regions (apex, middle and base). The observed gradient distribution of various subtypes along frequency axis suggest that these SGNs play different but complementing roles in processing various sound frequencies. Particularly, subtype I_a_ and I_b_ SGNs dominate in the low frequencies, whereas subtype I_c_ SGNs reciprocally contribute more in high frequencies. Such frequency-dependent distribution of SGN subtypes are consistent with previous studies in cats^16^ and gerbils^5, 20^ by sampling auditory nerve fibers using single unit recording, all of which reported higher percentage of HSR fibers in low frequencies and higher percentage of LSR fibers in high frequencies. Similar conclusions were also suggested by studies that analyzed the fiber diameter of the cochlear nerve in mice^17^ and monkeys^29^, which showed that fibers from the base turn have thinner diameter on average than those from the apical turn. Since LSR SGNs have thinner fiber diameter than HSR SGNs^2, 11^, these studies support a relatively higher percentage of LSR SGNs in the high frequency region. Consistently, the gradient distribution pattern of different SGNs was also observed at the gene expression level from single-cell RNA sequencing of SGNs^18^.

The observed loss of subtype I_c_ SGNs in middle-age and old mice agrees with the idea that LSR neurons are more vulnerable^19^ and are preferentially damaged during aging^27, 30, 31^. These SGNs have high threshold and are critical in the processing of loud signals, especially in noisy environment. Selective loss of these neurons, or even just the loss of their peripheral synapses with intact cell bodies (termed cochlear synaptopathy), is believed to underlie the phenomena of hidden hearing loss^32, 33^, under which the hearing threshold remains normal in quiet condition but signal detection in noisy environment is compromised. Remarkably, we found that the preferential loss of I_c_ SGNs during aging is progressively more profound in the base region of the cochlea (Fig. 3A). Since ARHL usually starts from the high frequency region and progresses toward the low frequency region^34, 35^, which occurs in parallel with the gradient loss of I_c_ SGNs along the frequency axis over time (Fig. 3A), it suggests that the more profound loss of I_c_ SGNs in the base region may uniquely drive the high frequency hearing loss during aging.

Despite the widely observed vulnerability in LSR SGNs under pathological conditions^19, 30, 36, 37^, the underlying mechanisms of their preferential damage remains unclear. One possible explanation may lie in their distinct molecular identity of not expressing CR and CB, two major calcium binding proteins in regulating intracellular calcium^38, 39^. Calcium is a ubiquitous intracellular messenger that controls crucial cellular function including life and death. The concentration of calcium is tightly regulated through various pathways, with high calcium being toxic under pathological conditions^40, 41^. CR and CB are among the most abundant calcium binding proteins in the brain that chelate free calcium and maintain intracellular calcium at a low level, which is protective. Lacking CR and CB may impose additional risk to subtype I_c_ SGNs under challenging conditions, leading to the preferential damage of synapses^27^ and eventually cell death^19, 30, 36, 37^.

One limitation of this study was that the classification of SGN subtypes was based on the expression patterns of only three molecular markers, including Tuj1, CR and CB. These markers have been recognized to label all type I, subtype I_a_ and subtype I_b_ SGNs^18, 21, 26^, respectively. Due to the technical difficulty we had with the Pou4f1 antibody in the staining, we used the Tuj1 antibody instead to identify all type I SGNs, and classified the cells without CR and CB expression as subtype I_c_ SGNs. These CR^-^ CB^-^ neurons are expected to express Pou4f1, but may not represent all Pou4f1-expressing SGNs due to the fact that some neurons express more than one of these markers^18, 21^. Indeed, we found many SGNs that express both CR and CB and analyzed these neurons in the dual-labeled I_a_/I_b_ group. Nonetheless, SGNs are far more heterogenous in their molecular identity beyond these selected markers^18, 21, 22^, and should not be considered as a simple collection of exclusively distinct subpopulations of neurons. The classification of four SGN subpopulations in this study is by no means an ideal way to unambiguously identify functionally distinct SGN subtypes, but rather a simplified approach to provide a convenient and valid method to systematically characterize the general distribution patterns of different SGNs and their age-related changes.

## Data availability statement

The original contributions presented in the study are included in the article, further inquiries can be directed to the corresponding author.

## Ethics statement

The animal study was reviewed and approved by the Institutional Animal Care and Use Committee of The Ohio State University.

## Author contributions

RX designed the study. MW, SL and RX collected the data. MW and RX analyzed the data. RX drafted the manuscript. All authors edited and approved the submitted version.

## References

1. Spoendlin H. Innervation patterns in the organ of corti of the cat. Acta Otolaryngol. 1969;67(2):239–54. Epub 1969/02/01. doi: 10.3109/00016486909125448. PubMed PMID: 5374642.

2. Liberman MC. Single-neuron labeling in the cat auditory nerve. Science. 1982;216(4551):1239–41. Epub 1982/06/11. doi: 10.1126/science.7079757. PubMed PMID: 7079757.

3. Liberman MC. Auditory-nerve response from cats raised in a low-noise chamber. The Journal of the Acoustical Society of America. 1978;63(2):442–55. Epub 1978/02/01. doi: 10.1121/1.381736. PubMed PMID: 670542.

4. Winter IM, Robertson D, Yates GK. Diversity of characteristic frequency rate-intensity functions in guinea pig auditory nerve fibres. Hear Res. 1990;45(3):191–202. Epub 1990/05/01. doi: 10.1016/0378-5955(90)90120-e. PubMed PMID: 2358413.

5. Schmiedt RA. Spontaneous rates, thresholds and tuning of auditory-nerve fibers in the gerbil: comparisons to cat data. Hear Res. 1989;42(1):23–35. Epub 1989/10/01. doi: 10.1016/0378-5955(89)90115-9. PubMed PMID: 2584157.

6. Borg E, Engstrom B, Linde G, Marklund K. Eighth nerve fiber firing features in normal-hearing rabbits. Hear Res. 1988;36(2-3):191–201. Epub 1988/11/01. doi: 10.1016/0378-5955(88)90061-5. PubMed PMID: 3209492.

7. Schalk TB, Sachs MB. Nonlinearities in auditory-nerve fiber responses to bandlimited noise. The Journal of the Acoustical Society of America. 1980;67(3):903–13. Epub 1980/03/01. doi: 10.1121/1.383970. PubMed PMID: 7358915.

8. Zeng FG, Turner CW, Relkin EM. Recovery from prior stimulation. II: Effects upon intensity discrimination. Hear Res. 1991;55(2):223–30. Epub 1991/10/01. doi: 10.1016/0378-5955(91)90107-k. PubMed PMID: 1757290.

9. Young ED, Barta PE. Rate responses of auditory nerve fibers to tones in noise near masked threshold. The Journal of the Acoustical Society of America. 1986;79(2):426–42. Epub 1986/02/01. doi: 10.1121/1.393530. PubMed PMID: 3950195.

10. Sachs MB, Voigt HF, Young ED. Auditory nerve representation of vowels in background noise. J Neurophysiol. 1983;50(1):27–45. Epub 1983/07/01. doi: 10.1152/jn.1983.50.1.27. PubMed PMID: 6875649.

11. Gleich O, Wilson S. The diameters of guinea pig auditory nerve fibres: distribution and correlation with spontaneous rate. Hear Res. 1993;71(1-2):69–79. Epub 1993/12/01. doi: 10.1016/0378-5955(93)90022-s. PubMed PMID: 7509334.

12. Liberman LD, Wang H, Liberman MC. Opposing gradients of ribbon size and AMPA receptor expression underlie sensitivity differences among cochlear-nerve/hair-cell synapses. J Neurosci. 2011;31(3):801–8. Epub 2011/01/21. doi: 10.1523/JNEUROSCI.3389-10.2011. PubMed PMID: 21248103; PMCID: PMC3290333.

13. Fekete DM, Rouiller EM, Liberman MC, Ryugo DK. The central projections of intracellularly labeled auditory nerve fibers in cats. J Comp Neurol. 1984;229(3):432–50. Epub 1984/11/01. doi: 10.1002/cne.902290311. PubMed PMID: 6209306.

14. Rouiller EM, Cronin-Schreiber R, Fekete DM, Ryugo DK. The central projections of intracellularly labeled auditory nerve fibers in cats: an analysis of terminal morphology. J Comp Neurol. 1986;249(2):261–78. Epub 1986/07/08. doi: 10.1002/cne.902490210. PubMed PMID: 3734159.

15. Liberman MC. Central projections of auditory-nerve fibers of differing spontaneous rate. I. Anteroventral cochlear nucleus. J Comp Neurol. 1991;313(2):240–58. Epub 1991/11/08. doi: 10.1002/cne.903130205. PubMed PMID: 1722487.

16. Liberman MC. Physiology of cochlear efferent and afferent neurons: direct comparisons in the same animal. Hear Res. 1988;34(2):179–91. Epub 1988/07/15. doi: 10.1016/0378-5955(88)90105-0. PubMed PMID: 3170360.

17. Anniko M, Arnesen AR. Cochlear nerve topography and fiber spectrum in the pigmented mouse. Archives of oto-rhino-laryngology. 1988;245(3):155–9. Epub 1988/01/01. doi: 10.1007/BF00464018. PubMed PMID: 3178564.

18. Shrestha BR, Chia C, Wu L, Kujawa SG, Liberman MC, Goodrich LV. Sensory Neuron Diversity in the Inner Ear Is Shaped by Activity. Cell. 2018;174(5):1229–46 e17. Epub 2018/08/07. doi: 10.1016/j.cell.2018.07.007. PubMed PMID: 30078709; PMCID: PMC6150604.

19. Furman AC, Kujawa SG, Liberman MC. Noise-induced cochlear neuropathy is selective for fibers with low spontaneous rates. J Neurophysiol. 2013;110(3):577–86. Epub 2013/04/19. doi: 10.1152/jn.00164.2013. PubMed PMID: 23596328; PMCID: PMC3742994.

20. Schmiedt RA, Mills JH, Boettcher FA. Age-related loss of activity of auditory-nerve fibers. J Neurophysiol. 1996;76(4):2799–803. Epub 1996/10/01. doi: 10.1152/jn.1996.76.4.2799. PubMed PMID: 8899648.

21. Sun S, Babola T, Pregernig G, So KS, Nguyen M, Su SM, Palermo AT, Bergles DE, Burns JC, Muller U. Hair Cell Mechanotransduction Regulates Spontaneous Activity and Spiral Ganglion Subtype Specification in the Auditory System. Cell. 2018;174(5):1247–63 e15. Epub 2018/08/07. doi: 10.1016/j.cell.2018.07.008. PubMed PMID: 30078710; PMCID: PMC6429032.

22. Petitpre C, Wu H, Sharma A, Tokarska A, Fontanet P, Wang Y, Helmbacher F, Yackle K, Silberberg G, Hadjab S, Lallemend F. Neuronal heterogeneity and stereotyped connectivity in the auditory afferent system. Nature communications. 2018;9(1):3691. Epub 2018/09/14. doi: 10.1038/s41467-018-06033-3. PubMed PMID: 30209249; PMCID: PMC6135759.

23. Sherrill HE, Jean P, Driver EC, Sanders TR, Fitzgerald TS, Moser T, Kelley MW. Pou4f1 Defines a Subgroup of Type I Spiral Ganglion Neurons and Is Necessary for Normal Inner Hair Cell Presynaptic Ca(2+) Signaling. J Neurosci. 2019;39(27):5284–98. Epub 2019/05/16. doi: 10.1523/JNEUROSCI.2728-18.2019. PubMed PMID: 31085606; PMCID: PMC6607758.

24. Sharma K, Seo YW, Yi E. Differential Expression of Ca(2+)-buffering Protein Calretinin in Cochlear Afferent Fibers: A Possible Link to Vulnerability to Traumatic Noise. Exp Neurobiol. 2018;27(5):397–407. Epub 2018/11/16. doi: 10.5607/en.2018.27.5.397. PubMed PMID: 30429649; PMCID: PMC6221833.

25. Bottom RT, Siebald C, Vincent PFY, Sun S, Reijntjes DOJ, Manca M, Glowatzki E, Muler U. Molecular signatures define subtypes of auditory afferent neurons with distinct peripheral projection patterns and physiological properties. ARO 2023 poster presentation #SA23. 2023.

26. Nishimura K, Noda T, Dabdoub A. Dynamic Expression of Sox2, Gata3, and Prox1 during Primary Auditory Neuron Development in the Mammalian Cochlea. PLoS One. 2017;12(1):e0170568. Epub 2017/01/25. doi: 10.1371/journal.pone.0170568. PubMed PMID: 28118374; PMCID: PMC5261741.

27. Wang M, Zhang C, Lin S, Wang Y, Seicol BJ, Ariss RW, Xie R. Biased auditory nerve central synaptopathy is associated with age-related hearing loss. J Physiol. 2021;599(6):1833–54. Epub 2021/01/16. doi: 10.1113/JP281014. PubMed PMID: 33450070; PMCID: PMC8197675.

28. Xie R. Transmission of auditory sensory information decreases in rate and temporal precision at the endbulb of Held synapse during age-related hearing loss. J Neurophysiol. 2016;116(6):2695–705. Epub 2016/09/30. doi: 10.1152/jn.00472.2016. PubMed PMID: 27683884; PMCID: PMC5133313.

29. Alving BM, Cowan WM. Some quantitative observations on the cochlear division of the eighth nerve in the squirrel monkey (Saimiri sciureus). Brain Res. 1971;25(2):229–39. Epub 1971/01/22. doi: 10.1016/0006-8993(71)90435-5. PubMed PMID: 4993585.

30. McClaskey CM, Dias JW, Schmiedt RA, Dubno JR, Harris KC. Evidence for Loss of Activity in Low-Spontaneous-Rate Auditory Nerve Fibers of Older Adults. J Assoc Res Otolaryngol. 2022;23(2):273–84. Epub 2022/01/13. doi: 10.1007/s10162-021-00827-x. PubMed PMID: 35020090; PMCID: PMC8964899.

31. Sergeyenko Y, Lall K, Liberman MC, Kujawa SG. Age-related cochlear synaptopathy: an early-onset contributor to auditory functional decline. J Neurosci. 2013;33(34):13686–94. Epub 2013/08/24. doi: 10.1523/JNEUROSCI.1783-13.2013. PubMed PMID: 23966690; PMCID: PMC3755715.

32. Plack CJ, Barker D, Prendergast G. Perceptual consequences of “hidden” hearing loss. Trends in hearing. 2014;18. Epub 2014/09/11. doi: 10.1177/2331216514550621. PubMed PMID: 25204468; PMCID: PMC4227662.

33. Liberman MC. Noise-induced and age-related hearing loss: new perspectives and potential therapies. F1000Res. 2017;6:927. Epub 2017/07/12. doi: 10.12688/f1000research.11310.1. PubMed PMID: 28690836; PMCID: PMC5482333.

34. Ison JR, Allen PD, O’Neill WE. Age-related hearing loss in C57BL/6J mice has both frequency-specific and non-frequency-specific components that produce a hyperacusis-like exaggeration of the acoustic startle reflex. J Assoc Res Otolaryngol. 2007;8(4):539–50. Epub 2007/10/24. doi: 10.1007/s10162-007-0098-3. PubMed PMID: 17952509; PMCID: PMC2538342.

35. Fetoni AR, Picciotti PM, Paludetti G, Troiani D. Pathogenesis of presbycusis in animal models: a review. Exp Gerontol. 2011;46(6):413–25. Epub 2011/01/08. doi: 10.1016/j.exger.2010.12.003. PubMed PMID: 21211561.

36. Liberman LD, Suzuki J, Liberman MC. Dynamics of cochlear synaptopathy after acoustic overexposure. J Assoc Res Otolaryngol. 2015;16(2):205–19. Epub 2015/02/14. doi: 10.1007/s10162-015-0510-3. PubMed PMID: 25676132; PMCID: PMC4368657.

37. Kujawa SG, Liberman MC. Synaptopathy in the noise-exposed and aging cochlea: Primary neural degeneration in acquired sensorineural hearing loss. Hear Res. 2015;330(Pt B):191–9. Epub 2015/03/15. doi: 10.1016/j.heares.2015.02.009. PubMed PMID: 25769437; PMCID: PMC4567542.

38. Schwaller B. Cytosolic Ca2+ buffers. Cold Spring Harbor perspectives in biology. 2010;2(11):a004051. Epub 2010/10/15. doi: 10.1101/cshperspect.a004051. PubMed PMID: 20943758; PMCID: PMC2964180.

39. Rogers J, Khan M, Ellis J. Calretinin and other CaBPs in the nervous system. Adv Exp Med Biol. 1990;269:195–203. Epub 1990/01/01. doi: 10.1007/978-1-4684-5754-4_32. PubMed PMID: 2191557.

40. Foster TC. Calcium homeostasis and modulation of synaptic plasticity in the aged brain. Aging cell. 2007;6(3):319–25. Epub 2007/05/23. doi: 10.1111/j.1474-9726.2007.00283.x. PubMed PMID: 17517041.

41. Berridge MJ. Calcium signalling remodelling and disease. Biochemical Society transactions. 2012;40(2):297–309. Epub 2012/03/23. doi: 10.1042/BST20110766. PubMed PMID: 22435804.

